# BGC-QUAST: a quality assessment tool for genome mining software

**DOI:** 10.64898/2026.05.04.722653

**Authors:** Aleksandra Kushnareva, Daria Tupikina, Hesham Almessady, Alice C. McHardy, Alexey Gurevich

## Abstract

**Summary:** Biosynthetic gene clusters (BGCs) encode microbial natural products, many of which have important ecological and biomedical roles. Genome mining tools enable large-scale BGC prediction, but their outputs differ substantially, complicating comparison and interpretation. We present BGC-QUAST, a framework for evaluating and comparing BGC predictions across three analysis modes: comparison across samples, assessment of BGC recovery in draft assemblies relative to reference genomes, and comparison of predictions from different tools using overlap analysis. BGC-QUAST provides standardized metrics, interactive visualizations, and integrated outputs for joint inspection of predictions, enabling the comprehensive comparison of genome mining results and facilitating sample prioritisation based on biosynthetic potential.

**Availability and implementation:** BGC-QUAST is publicly available at https://github.com/gurevichlab/bgc-quast

## 1 Introduction

Microbial natural products represent a major source of therapeutic agents, including clinically important antibiotics (Newman and Cragg 2020). These compounds are typically encoded in biosynthetic gene clusters (BGCs), functional groups of physically clustered genes. Advances in next-generation sequencing have increased the availability of genomic data, enabling the direct identification of BGCs from genome sequences, i.e., genome mining, leading to the development of a range of computational methods for BGC detection (Biermann et al. 2022).

Representative BGC prediction tools include rule- and profile-based methods such as antiSMASH (Blin et al. 2025) and PRISM (Skinnider et al. 2020), as well as machine-learning approaches such as DeepBGC (Hannigan et al. 2019) and GECCO (Carroll et al. 2021). Complementary query-guided tools, such as cblaster (Gilchrist et al. 2021) and GATOR-GC (Cediel-Becerra et al. 2025), support targeted genome mining by identifying and comparing BGCs related to a selected query gene, protein, or BGC.

Despite the growing number of BGC prediction tools, systematic and reproducible comparison of their results remains difficult. First, benchmarking is restricted by the lack of comprehensive ground truth datasets. The most extensive curated resource, the Minimum Information about a Biosynthetic Gene cluster (MIBiG) database (Zdouc et al. 2025), contains a limited number of experimentally validated BGCs and is widely used as training and testing data, reducing its suitability for benchmarking. In addition, outputs from different tools vary in both format and content. Differences include file structure, BGC product nomenclature, and cluster boundary and completeness definitions.

As a result, predictions generated for the same genome by different methods can vary substantially in length and gene content. Although existing workflow frameworks, such as nf-core/funcscan (Fellows Yates et al. 2026), can integrate multiple genome mining tools and aggregate their outputs, they do not provide standardized quality metrics for evaluating and comparing BGC predictions across tools and datasets (Ewels et al. 2020).

Here, we present BGC-QUAST, a tool for quality assessment and comparative analysis of BGC predictions. It supports three complementary analysis modes: comparison across samples, evaluation of assembly-based predictions against high-quality reference genomes, and comparison of output from different genome mining tools. BGC-QUAST is intended for both developers and users of genome mining methods, enabling quantitative comparison of predictions and generating interactive reports along with publication-ready summaries and visualizations.

## 2 Methods

### 2.1 Input data and preprocessing

An overview of the BGC-QUAST workflow is shown in Fig. 1. As primary input, BGC-QUAST takes BGC predictions generated by genome mining tools. We currently support antiSMASH, DeepBGC, GECCO, and PRISM. Outputs from other genome mining tools can also be processed if formatted according to one of the supported tool schemas, as described in the BGC-QUAST documentation. For the compare-to-reference mode, assembly-to-reference coordinate mappings are required and are derived from QUAST output (Mikheenko et al. 2023). Genome sequences in FASTA or GenBank format can be provided for additional information in the report.

**Figure 1.**
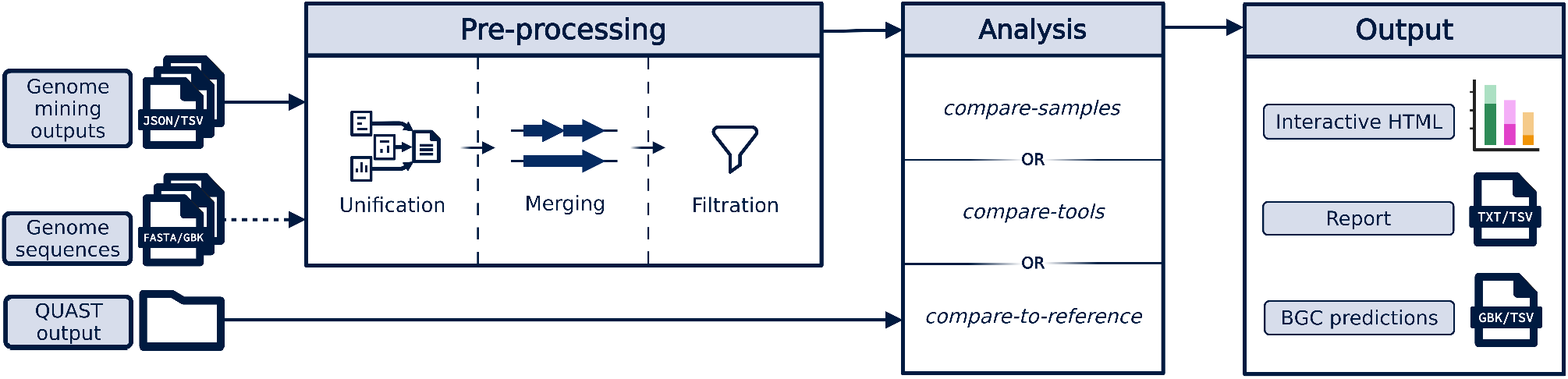
BGC-QUAST pipeline overview. BGC predictions from genome mining tools are converted into a unified format, optionally merged and filtered by user-specified length. Depending on the selected analysis mode, predictions are compared across genomes/samples, against a reference genome using QUAST alignments, or across tools, and summarized in an interactive report together with tabular outputs and files for downstream analysis.

Before analysis, all input predictions are converted into a unified internal format, where each BGC is defined by its genomic coordinates, predicted product type, completeness, and associated metadata. For consistency, tool-specific product type labels are mapped to a set of predefined product categories. BGC completeness is determined by the distance between predicted cluster boundaries and contig edges; a BGC is considered fragmented if either boundary lies within a specified distance from a contig edge (--edge-distance, default 100 bp), and complete otherwise.

Optional preprocessing steps include filtering BGCs by length (--min-length, default 0 bp) and merging adjacent BGC regions (--merge-distance, default 0 bp).

### 2.2 Analysis modes

Across all modes, quality metrics are reported both overall and, where applicable, stratified by BGC product type and completeness. The compare-tools mode operates on predictions from multiple genome mining tools applied to the same genome sequence. In contrast, the other two modes operate on predictions from the same genome mining tool applied to different genome sequences.

#### 2.2.1 Compare-samples mode

This mode summarizes and compares biosynthetic potential across multiple genomes or metagenomic samples. It serves as the basic analysis mode of BGC-QUAST, meaning that all reported metrics are also available in other modes.

The following metrics are computed for each input:

- **BGC counts**. Total number of predicted BGCs.
- **Length statistics**. Mean and total BGC length.
- **Gene count statistics**. If genome annotation is provided, the mean number of genes per BGC.

#### 2.2.2 Compare-to-reference mode

This mode assesses how well BGCs predicted on draft assemblies, including metagenome-assembled genomes (MAGs), match those in a high-quality reference genome, thereby evaluating the robustness of BGC detection to assembly fragmentation.

Assembly-to-reference alignments are obtained from QUAST output. BGC-QUAST then maps BGCs predicted in draft assemblies to reference BGCs using these alignments. Based on this comparison, the following metrics are reported per input assembly and, where applicable, for the reference genome:

- **Fully recovered BGCs**. Reference BGCs with ≥95% of their length covered by one (single-contig) or multiple (multi-contig) assembly BGCs.
- **Partially recovered BGCs**. Reference BGCs with 10–95% coverage by one or more assembly BGCs.
- **Missed BGCs**. Reference BGCs with <10% coverage.
- **Recovery rate**. Proportion of reference BGCs that are fully or partially recovered.
- **Product type mismatches**. Assembly BGCs mapped to reference BGCs but predicted with a different product type.
- **Unmapped assembly BGCs**. Assembly BGCs that do not map to any reference BGC.

#### 2.2.3 Compare-tools mode

This mode compares BGC predictions from different genome mining tools applied to the same genome sequence using coordinate-based overlap matching. Each predicted BGC is treated as a genomic interval defined by the start and end coordinates reported by the corresponding genome mining tool.

For each BGC predicted by a given tool, BGC-QUAST calculates the fraction of its interval overlapped by each BGC predicted by another tool. If at least one such overlap fraction reaches the defined threshold (--overlap-fraction, default 0.9, i.e., 90% of the BGC length), the BGC is considered shared with respect to that tool; otherwise, it is classified as unique (Supplementary Note S1). Overlaps are evaluated independently for each tool. For example, if a BGC predicted by one tool lies entirely within a larger BGC from another, the smaller BGC is classified as shared, while the larger one may still be considered unique if the overlap is below the threshold.

The following metrics are reported per tool:

- **Unique BGCs**. Number of BGCs uniquely identified by the tool. A BGC is considered unique overall if it is classified as unique with respect to all other tools.
- **Unique rate**. Proportion of BGCs detected exclusively by the tool relative to its total number of predictions.

Pairwise overlaps between tools are visualized using Venn diagrams to show the number of unique and shared BGCs. Two values are reported in the intersection of the Venn diagram, reflecting asymmetric counts of shared BGCs computed separately for each tool.

In addition, BGC-QUAST generates a combined GenBank file containing all predictions from the selected tools in a unified coordinate framework, as well as tab-separated (TSV) outputs summarizing all predicted BGCs and grouping overlapping predictions into genomic intervals for side-by-side comparison, enabling manual inspection and downstream analysis.

## 3 Results

To demonstrate the utility of BGC-QUAST across different use cases, we evaluated the tool in all three analysis modes using reference genomes and MAGs from the CAMI II plant-associated dataset (Meyer et al. 2022). The analyzed datasets are listed in Supplementary Table S1. Runtime and peak RAM measurements for the genome mining tools, QUAST, and BGC-QUAST are summarized in Supplementary Table S2.

### 3.1 Compare-samples mode identifies genomes with high biosynthetic potential

We first applied BGC-QUAST in the compare-samples mode to antiSMASH genome mining results obtained for a small set of reference genomes. The selected ten genomes represent diverse orders within three bacterial phyla known for their high biosynthetic potential: Actinomycetota, Cyanobacteriota, and Myxococcota (Supplementary Table S1).

Across these genomes, BGC-QUAST revealed substantial variation in both the number and mean length of predicted biosynthetic gene clusters (BGCs). In particular, *Mycolicibacterium neoaurum* (PRJNA177066) exhibited the highest biosynthetic potential, with 21 predicted BGCs, compared to 5–15 BGCs across the remaining genomes. Mean BGC lengths varied nearly two-fold, from 23.5 kbp for *Jiangella gansuensis* (PRJNA63165) to 42.3 kbp for *Actinoplanes friuliensis* (PRJNA178199) (Supplementary Fig. S1).

As all input genomes are high-quality references, all identified BGCs were complete. At the same time, BGC-QUAST highlighted substantial variation in predicted BGC product types, with some genomes lacking specific classes; for example, *Pseudarthrobacter chlorophenolicus* (PRJNA20011) contains no ribosomally synthesized and post-translationally modified peptide (RiPP) or terpene BGCs, while *Thermobispora bispora* (PRJNA20737) lacks nonribosomal peptide synthetase (NRPS) BGCs (Supplementary Fig. S1).

These results demonstrate that BGC-QUAST can effectively rank genome sequences by biosynthetic potential while providing detailed insights beyond simple BGC counts, facilitating the selection of the most promising genomes or samples for downstream analyses.

### 3.2 Assembly quality strongly affects BGC recovery

Next, we used the compare-to-reference mode to quantify the impact of assembly quality on BGC recovery. We selected the reference genome with the highest number of BGCs from the compare-samples analysis (*M. neoaurum*) and compared it with the corresponding MAGs reconstructed using short-read (Otu207-short) and hybrid (Otu207-hybrid) gold-standard assemblies (Meyer et al. 2022).

Despite the high genome fraction of both MAGs (96.6% for Otu207-short and 99.4% for Otu207-hybrid), the quality of BGC predictions differs substantially. The number of predicted BGCs in the reference genome (21) is exceeded in Otu207-hybrid (23) and further increased in Otu207-short (29). However, this apparent increase does not reflect the detection of additional BGCs, but rather the extensive fragmentation of long clusters. This is evident from the mean BGC length, which decreases from 37.4 kbp in the reference genome to 22.9 kbp in Otu207-hybrid and just 5.7 kbp in Otu207-short. Consistently, stratification by completeness shows that Otu207-short contains no single complete BGCs, while Otu207-hybrid has only six complete clusters out of 23 (Supplementary Fig. S2).

Overall, the hybrid-based MAG fails to recover one reference BGC (recovery rate 95.2%), whereas the short-read-based MAG misses six BGCs (recovery rate 71.4%). When considering only fully recovered BGCs assembled in a single contig, Otu207-hybrid contains six such clusters, while Otu207-short contains none, with all recovered BGCs being partial.

These findings highlight the sensitivity of BGC detection to assembly contiguity and show that hybrid assembly substantially improves BGC recovery compared to the short-read-only approach, but still falls short of recovering complete BGCs present in the reference genome. At the same time, they illustrate how BGC-QUAST quantifies BGC recovery and fragmentation, enabling direct assessment of assembly quality in the context of genome mining.

### 3.3 Different genome mining tools produce complementary BGC predictions

Finally, we applied BGC-QUAST in the compare-tools mode to assess differences between BGC predictions made with antiSMASH, DeepBGC, and GECCO on the selected reference genome (*M. neoaurum*). PRISM was not included in this comparison as it is currently available only as a web service and could therefore not be evaluated under the same local computational setup as the other tools (Supplementary Table S2). The three compared tools demonstrated substantial variation in the number and characteristics of predicted BGCs (Supplementary Fig. S3).

In particular, DeepBGC predicts a markedly higher number of BGCs (97) compared to antiSMASH (21) and GECCO (12). However, more than half of DeepBGC predictions (56 out of 97) are assigned to the “unknown” product class, reflecting the uncertainty of the machine learning–based approach. In contrast, antiSMASH, which relies on curated rule- and HMM-based models, does not produce such “unknown” predictions by design.

The tools also differ greatly in predicted BGC lengths. antiSMASH reports considerably longer clusters (mean length 37.4 kbp) compared to GECCO (19.4 kbp) and DeepBGC (16.8 kbp), consistent with its conservative boundary detection strategy (Blin et al. 2025).

Despite the large number of predictions by DeepBGC, each tool identifies a distinct set of unique BGCs. antiSMASH retains 16 unique clusters (76% of its predictions), whereas GECCO identifies only a single unique BGC (Supplementary Fig. S3). These differences are further illustrated by pairwise overlap analyses (Supplementary Fig. S4), where Venn diagrams reveal limited agreement between tools and asymmetric overlaps arising from differences in predicted BGC lengths and boundaries.

These results highlight the ability of BGC-QUAST to systematically compare genome mining tools, revealing both shared and tool-specific BGC predictions. Such comparisons can support the evaluation of new prediction methods against established tools, the assessment of how different settings of the same tool affect its predictions, or the identification of methods that best recover experimentally characterized BGCs added, for example, in newer MIBiG releases. By combining quantitative metrics with pairwise overlap visualizations, tab-separated (TSV) outputs, an interactive HTML interval-based summary of BGC coordinates across tools (Supplementary Fig. S5), and a single GenBank file containing predictions from all tools, BGC-QUAST enables joint inspection of complementary predictions and provides a more complete view of an organism’s biosynthetic potential.

## 4 Discussion

BGC-QUAST provides a structured framework for summarizing and comparing BGC predictions across genomes, assemblies, and genome mining tools. It defines consistent metrics and organizes results in a unified and accessible format.

By design, BGC-QUAST does not assess the biological correctness of predicted BGCs, which ultimately requires experimental validation. Instead, it focuses on comparative evaluation of genome mining outputs, enabling identification of differences in predictions, recovery, and fragmentation across datasets and tools.

We expect BGC-QUAST to be useful for both users and developers of genome mining tools: it enables prioritization of genomes and samples based on biosynthetic potential, supports joint analysis of complementary predictions from multiple tools, and provides a framework for method comparison and refinement.

Future extensions of BGC-QUAST could include scalable grouping of related BGCs across samples using approaches such as BiG-SCAPE (Draisma et al. 2026), followed by more detailed gene-level inspection of related BGCs in the spirit of tools such as BisCEET (Sowa et al. 2026). Such functionality could facilitate the identification and prioritization of novel or sample-specific candidate BGCs for experimental validation. To further improve accessibility and promote wider adoption, we plan to extend BGC-QUAST with a web-based interface.

## Acknowledgements

We thank Lucy Ehrig for early contributions to this work, Judith Gadau for testing the tool, and Dr. Azat Tagirdzhanov for helpful discussions.

## Funding

This work has been funded by the German Research Foundation (DFG)—project number 460129525 (NFDI4Microbiota) and the German Center for Infection Research (TI 12.002) (H.A., A.C.M.), and by the Helmholtz UNLOCK Benchmarking call (A.K., A.G.).

### Conflict of Interest

none declared.

## Supplementary information

Supplementary data are available online. The genomes used in the Results section, the corresponding genome mining outputs, full BGC-QUAST reports, and the BGC-QUAST v1.1.0 release snapshot are available at Zenodo (DOI: 10.5281/zenodo.20025178). Commands and software versions used are in Supplementary Note S2.

## Notes

https://zenodo.org/records/20025178

### Competing Interest Statement

The authors have declared no competing interest.

### Summary of Updates

This version of the manuscript has been revised to clarify the scope of BGC-QUAST and its computational requirements in the manuscript and Supplementary Information. We also extended the software functionality and usability by adding support for PRISM, new compare-tools outputs in TSV and HTML formats, the option to choose antiSMASH annotation levels (region, candidate cluster, protocluster), simplified installation, and improved documentation. In addition, we fixed reported bugs and functional issues as well as manuscript deficiencies. The revised submission also includes an updated Figure 1, minor changes in affiliations, five new and one substituted references, a new Supplementary Table S2, a new Supplementary Figure S5, and a new Supplementary Note S1. The updated BGC-QUAST software has been released as version 1.1.0, and the corresponding reports were regenerated with the new version.

