## Supplementary data for "BGC-QUAST: a quality assessment tool for genome mining software"

[1]Helmholtz Institute for Pharmaceutical Research Saarland (HIPS), Helmholtz Centre for Infection Research (HZI), Saarbrücken, Germany [2]Center for Bioinformatics, Saarland University, Saarbrücken, Germany [3]Computational Biology of Infection Research, Helmholtz Centre for Infection Research (HZI), Braunschweig, Germany [4]Braunschweig Integrated Centre of Systems Biology (BRICS), Technische Universität Braunschweig, Braunschweig, Germany, [5]German Center for Infection Research (DZIF), partner site Hannover-Braunschweig, Braunschweig, Germany, [6]Department of Computer Science, Saarland University, Saarbrücken, Germany, [7]PharmaScienceHub, Saarbrücken, Germany

[†] These authors contributed equally to this work.

**Supplementary Table S1.** Genome sequences used in the demonstration experiments. The table includes 10 reference genomes and two metagenome-assembled genomes (MAGs) from the CAMI II plant-associated dataset (https://cami-challenge.org/datasets/plant-associated/). Genome identifiers correspond to NCBI BioProject accessions for reference genomes. The two MAGs are denoted using CAMI naming for the ground-truth bin corresponding to the PRJNA177066 reference (Otu207), with suffixes indicating the underlying gold-standard assembly (GSA): "hybrid" for the GSA constructed from short-read Illumina and long-read Oxford Nanopore data, and "short" for the GSA constructed from short-read-only data. The "Running mode" column indicates the BGC-QUAST analysis modes in which each genome was used: S, compare-samples; R, compare-to-reference; T, compare-tools. Genome length and number of contigs are reported for sequences ≥1 kbp, corresponding to the default cutoff used in genome mining analyses.

| Genome ID | Running mode | Taxonomy | | | Length (Mbp) | Number of contigs |
|---|---|---|---|---|---|---|
| | | Organism (Strain) | Phylum | Order | | |
| PRJNA20011 | S | *Pseudarthrobacter chlorophenolicus* A6 | Actinomycetota | Micrococcaceae | 4.40 | 1 |
| PRJNA20737 | S | *Thermobispora bispora* DSM 43833 | Actinomycetota | Streptosporangiaceae | 4.19 | 1 |
| PRJNA21089 | S | *Kribbella flavida* DSM 17836 | Actinomycetota | Kribbellaceae | 7.58 | 1 |
| PRJNA29547 | S | *Geodermatophilus obscurus* DSM 43160 | Actinomycetota | Geodermatophilaceae | 5.32 | 1 |
| PRJNA63165 | S | *Jiangella gansuensis* DSM 44835 | Actinomycetota | Jiangellaceae | 5.59 | 1 |
| PRJNA177066 | S/R/T | *Mycolicibacterium neoaurum* VKM Ac-1815D | Actinomycetota | Mycobacteriaceae | 5.42 | 1 |
| PRJNA178199 | S | *Actinoplanes friuliensis* DSM 7358 | Actinomycetota | Micromonosporaceae | 9.38 | 1 |
| PRJNA9606 | S | *Gloeobacter violaceus* PCC 7421 | Cyanobacteriota | Gloeobacterales | 4.66 | 1 |
| PRJNA158839 | S | *Allocoleopsis franciscana* PCC 7113 | Cyanobacteriota | Coleofasciculales | 7.47 | 1 |
| PRJNA12634 | S | *Anaeromyxobacter dehalogenans* 2CP-C | Myxococcota | Cystobacterineae | 5.01 | 1 |
| Otu207-hybrid | R | *Mycolicibacterium neoaurum* VKM Ac-1815D | Actinomycetota | Mycobacteriaceae | 5.39 | 317 |
| Otu207-short | R | *Mycolicibacterium neoaurum* VKM Ac-1815D | Actinomycetota | Mycobacteriaceae | 5.24 | 1406 |

**Supplementary Table S2.** Runtime and peak RAM usage of genome mining tools, QUAST, and BGC-QUAST on the datasets used in this study. For computational resource benchmarking, we included genome mining tools that could be executed locally from the command line under the same computational setup. Measurements were performed on a Linux server node with 128 CPUs and 503 GiB of RAM. All programs were run sequentially using one CPU. Runtime (wall-clock time) and peak RAM usage (maximum resident set size) were recorded using GNU `/usr/bin/time -v`. Genome mining tools were benchmarked separately on each of the 10 reference genomes used in the compare-samples analysis and the two MAGs used in the compare-to-reference analysis. Results were summarized separately for the reference genomes and MAGs. For datasets analyzed in multiple separate runs, we report total runtime, mean ± standard deviation (SD) runtime, and the highest peak RAM across runs. QUAST was benchmarked in a single run using the reference genome and the two corresponding MAGs, and each BGC-QUAST analysis mode was benchmarked using the corresponding inputs. Mean ± SD did not apply to these single runs and is indicated by “n/a”.

| Software or analysis mode | Input dataset | Total runtime (h:mm:ss) | Mean ± SD runtime (mm:ss) | Peak RAM (MB) |
|---|---|---|---|---|
| **Generation of analysis inputs** | | | | |
| antiSMASH | 10 reference genomes | 0:13:23 | 01:20 ± 00:28 | 350.5 |
| antiSMASH | 2 MAGs | 0:09:59 | 05:00 ± 02:27 | 399.6 |
| GECCO | 10 reference genomes | 0:13:58 | 01:24 ± 00:23 | 1213.5 |
| GECCO | 2 MAGs | 0:02:34 | 01:17 ± 00:03 | 1111.7 |
| DeepBGC | 10 reference genomes | 3:56:47 | 23:41 ± 07:02 | 768.5 |
| DeepBGC | 2 MAGs | 0:18:32 | 09:16 ± 11:09 | 703.4 |
| QUAST | Reference genome and two MAGs | 0:00:27 | n/a | 151.9 |
| **BGC-QUAST analyses** | | | | |
| Compare-samples | antiSMASH outputs for 10 reference genomes | 0:00:11 | n/a | 136.5 |
| Compare-tools | antiSMASH, GECCO, and DeepBGC outputs for the reference PRJNA177066 | 0:00:08 | n/a | 120.0 |
| Compare-to-reference | antiSMASH outputs for the reference PRJNA177066 and two MAGs, plus QUAST output | 0:00:08 | n/a | 134.3 |

### Supplementary Figures

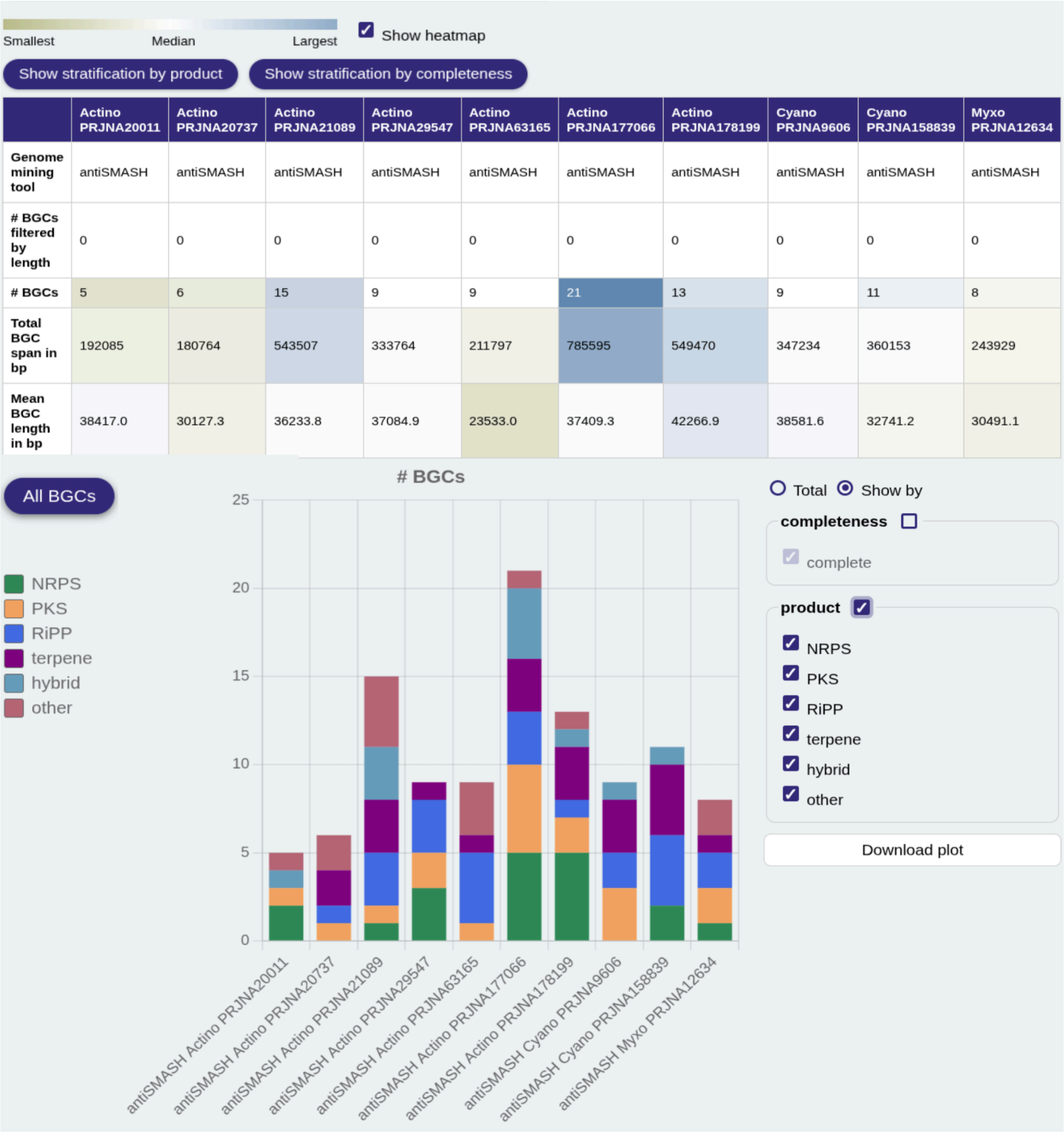

| | Actino PRJNA20011 | Actino PRJNA20737 | Actino PRJNA21089 | Actino PRJNA29547 | Actino PRJNA63165 | Actino PRJNA177066 | Actino PRJNA178199 | Cyano PRJNA9606 | Cyano PRJNA158839 | Myxo PRJNA12634 |
|---|---|---|---|---|---|---|---|---|---|---|
| **Genome mining tool** | antiSMASH | antiSMASH | antiSMASH | antiSMASH | antiSMASH | antiSMASH | antiSMASH | antiSMASH | antiSMASH | antiSMASH |
| **# BGCs filtered by length** | 0 | 0 | 0 | 0 | 0 | 0 | 0 | 0 | 0 | 0 |
| **# BGCs** | 5 | 6 | 15 | 9 | 9 | 21 | 13 | 9 | 11 | 8 |
| **Total BGC span in bp** | 192085 | 180764 | 543507 | 333764 | 211797 | 785595 | 549470 | 347234 | 360153 | 243929 |
| **Mean BGC length in bp** | 38417.0 | 30127.3 | 36233.8 | 37084.9 | 23533.0 | 37409.3 | 42266.9 | 38581.6 | 32741.2 | 30491.1 |



**Supplementary Figure S1**. Example fragment of a BGC-QUAST compare-samples report. The top panel summarizes, for each analyzed genome (7 actinomycetes, 2 cyanobacteria, and 1 myxobacterium from the CAMI II plant-associated dataset), the total number of predicted biosynthetic gene clusters (BGCs) and their mean length in base pairs (bp), as identified by antiSMASH. The heatmap highlights the minimum and maximum values for each metric across genomes. The controls above the table allow stratification of these metrics by BGC product type and completeness. The bottom panel provides a graphical representation of BGC counts across genomes as a stacked bar chart; here, counts are stratified by product type (colors; see legend), although alternative views (e.g., total counts) are also supported. This report enables identification of samples with the highest and lowest overall and class-specific biosynthetic potential.

Smallest Median Largest ☑ Show heatmap ☑ Include reference

Show stratification by product | Show stratification by completeness

| | reference | Otu207-hybrid | Otu207-short |
|---|---|---|---|
| **Genome mining tool** | antiSMASH | antiSMASH | antiSMASH |
| **# BGCs filtered by length** | 0 | 0 | 0 |
| **# BGCs** | 21 | 23 | 29 |
| **Total BGC span in bp** | 785595 | 526872 | 164477 |
| **Mean BGC length in bp** | 37409.3 | 22907.5 | 5671.6 |
| **# full ref. BGCs** | 21 | 6 | 0 |
| **# full ref. BGCs, single-contig** | 21 | 6 | 0 |
| **# full ref. BGCs, multi-contig** | 0 | 0 | 0 |
| **# partial ref. BGCs** | 0 | 14 | 15 |
| **# missed ref. BGCs** | 0 | 1 | 6 |
| **Ref. BGC recovery rate** | 1.00 | 0.95 | 0.71 |
| **# product type mismatches to ref.** | 0 | 0 | 0 |
| **# unmapped BGCs to ref.** | 0 | 0 | 0 |

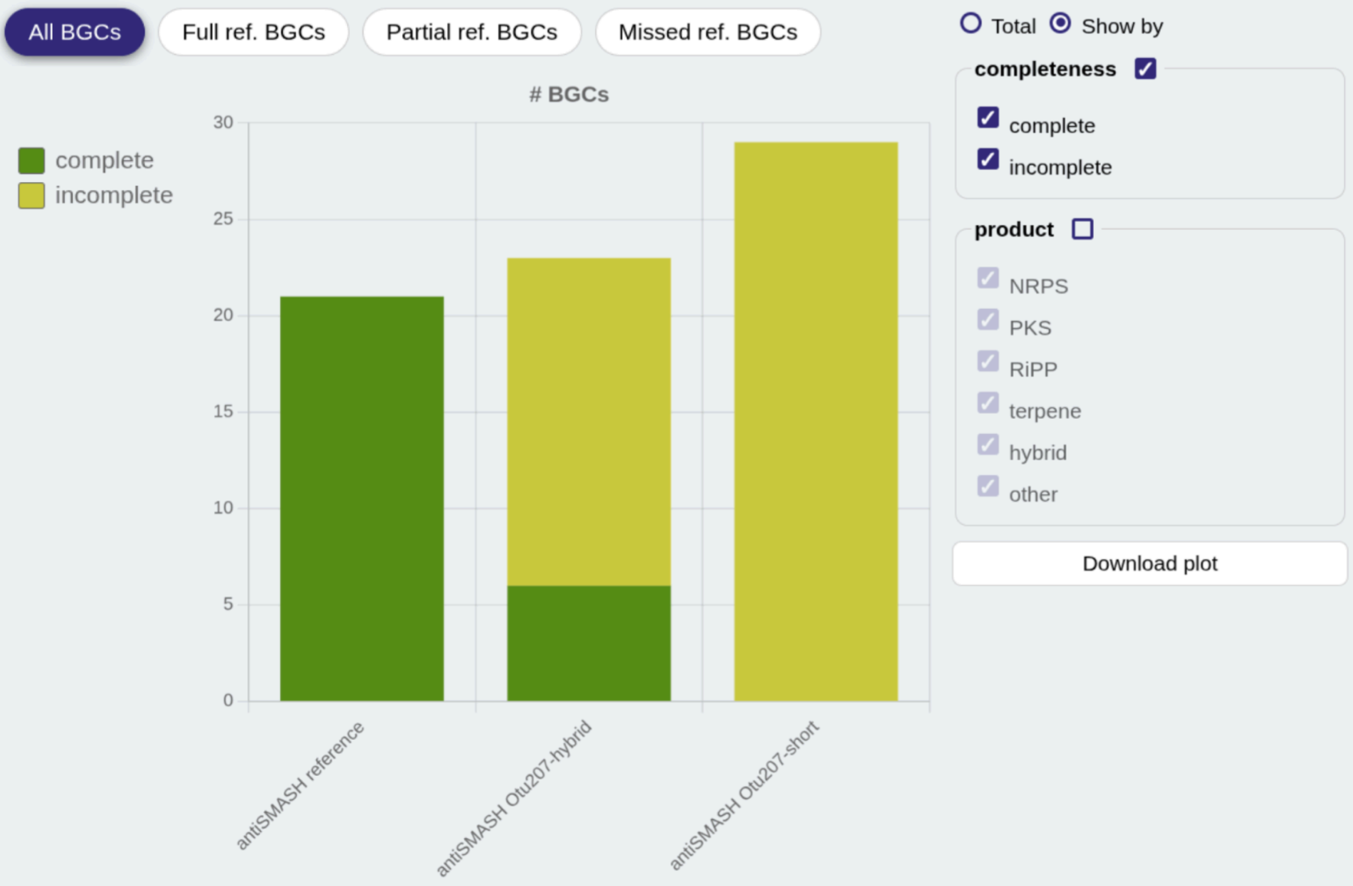


**Supplementary Figure S2**. Example fragment of a BGC-QUAST compare-to-reference report. The report compares biosynthetic gene clusters (BGCs) identified by antiSMASH in a reference genome and two corresponding metagenome-assembled genomes (MAGs) derived from gold-standard assemblies of the CAMI II plant-associated dataset, constructed using hybrid and short-read data. In this dataset, contigs are grouped into bins corresponding to individual reference genomes; here, both MAGs (Otu207) originate from the same reference genome, PRJNA177066 (*Mycolicibacterium neoaurum* VKM Ac-1815D). The top panel summarizes the total number of BGCs, their mean length in base pairs (bp), and their correspondence to BGCs predicted in the reference (first column). BGCs are categorized as fully recovered (in a single or multiple contigs), partially recovered, or missed with respect to the reference; the overall BGC recovery rate includes both fully and partially recovered BGCs. The bottom panel provides a graphical representation of BGC counts across samples, here stratified by completeness (complete, green; incomplete, yellow). This report enables assessment of how assembly quality impacts BGC reconstruction, as illustrated here by the short-read assembly yielding predominantly fragmented (incomplete) and shorter predicted BGCs compared to the hybrid assembly and, more markedly, the complete high-quality reference.

Smallest Median Largest

☑ Show heatmap

Show stratification by product | Show stratification by completeness

| | reference | reference | reference |
|---|---|---|---|
| **Genome mining tool** | antiSMASH | GECCO | DeepBGC |
| **# BGCs filtered by length** | 0 | 0 | 0 |
| **# BGCs** | 21 | 12 | 97 |
| **Total BGC span in bp** | 785595 | 233164 | 1631021 |
| **Mean BGC length in bp** | 37409.3 | 19430.3 | 16814.6 |
| **# unique BGCs** | 16 | 1 | 84 |
| **Unique BGC rate** | 0.76 | 0.08 | 0.87 |

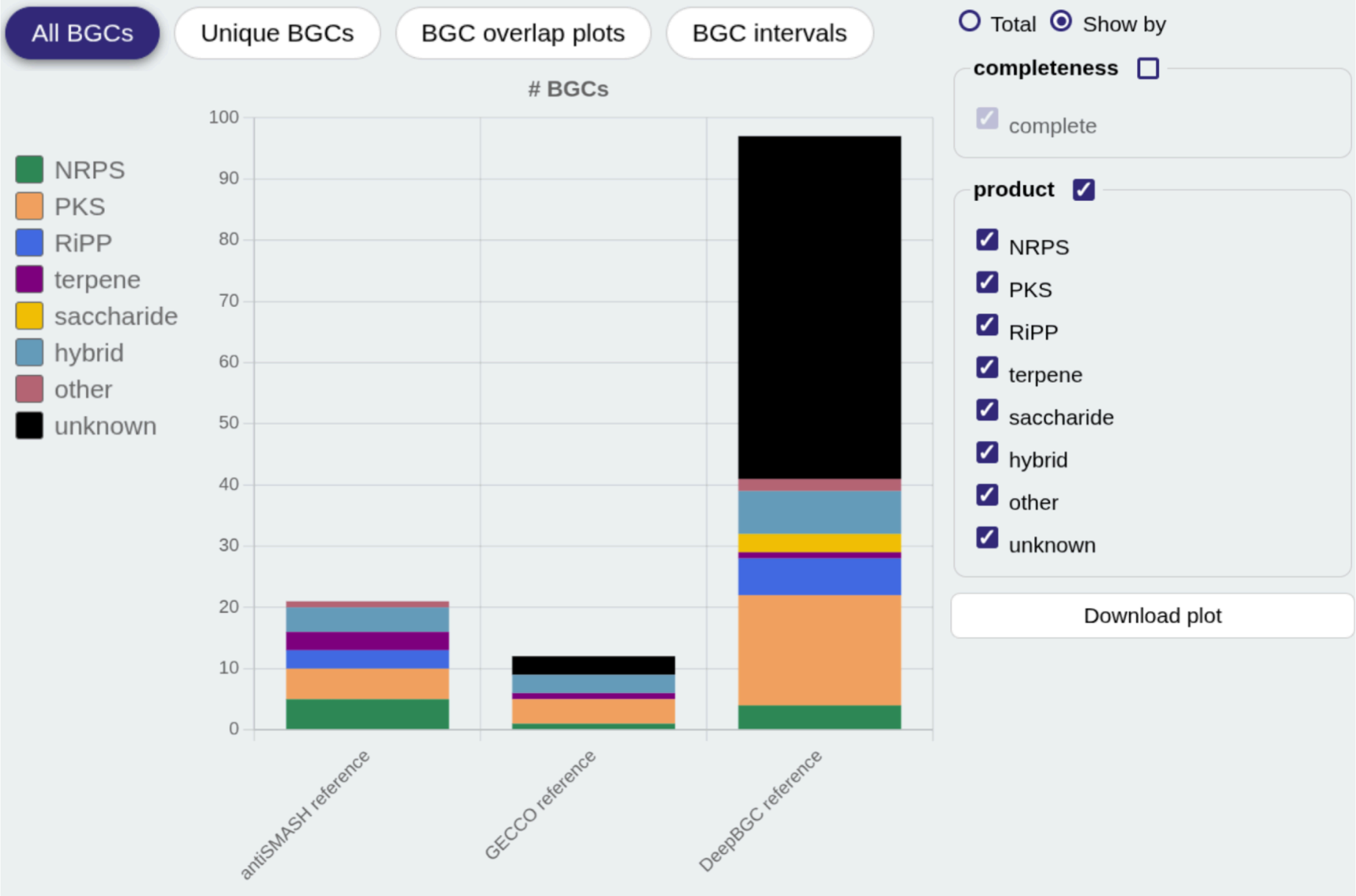


**Supplementary Figure S3.** Example fragment of a BGC-QUAST compare-tools report. The report compares biosynthetic gene clusters (BGCs) predicted in the same reference genome (here, PRJNA177066: *Mycolicibacterium neoaurum* VKM Ac-1815D) by three genome mining tools: antiSMASH, GECCO, and DeepBGC. The top panel summarizes the total number of predicted BGCs, their mean length in base pairs (bp), and the number and fraction of unique BGCs identified by each tool. A BGC is considered unique to a tool if no BGC predicted by another tool overlaps at least the defined fraction of its length (default threshold: 90%). The bottom panel provides a graphical representation of BGC counts across the different genome mining tools; here, counts are stratified by product type (colors; see legend). This report enables comparison of tool-specific sensitivity and agreement, highlighting differences in BGC detection across methods.

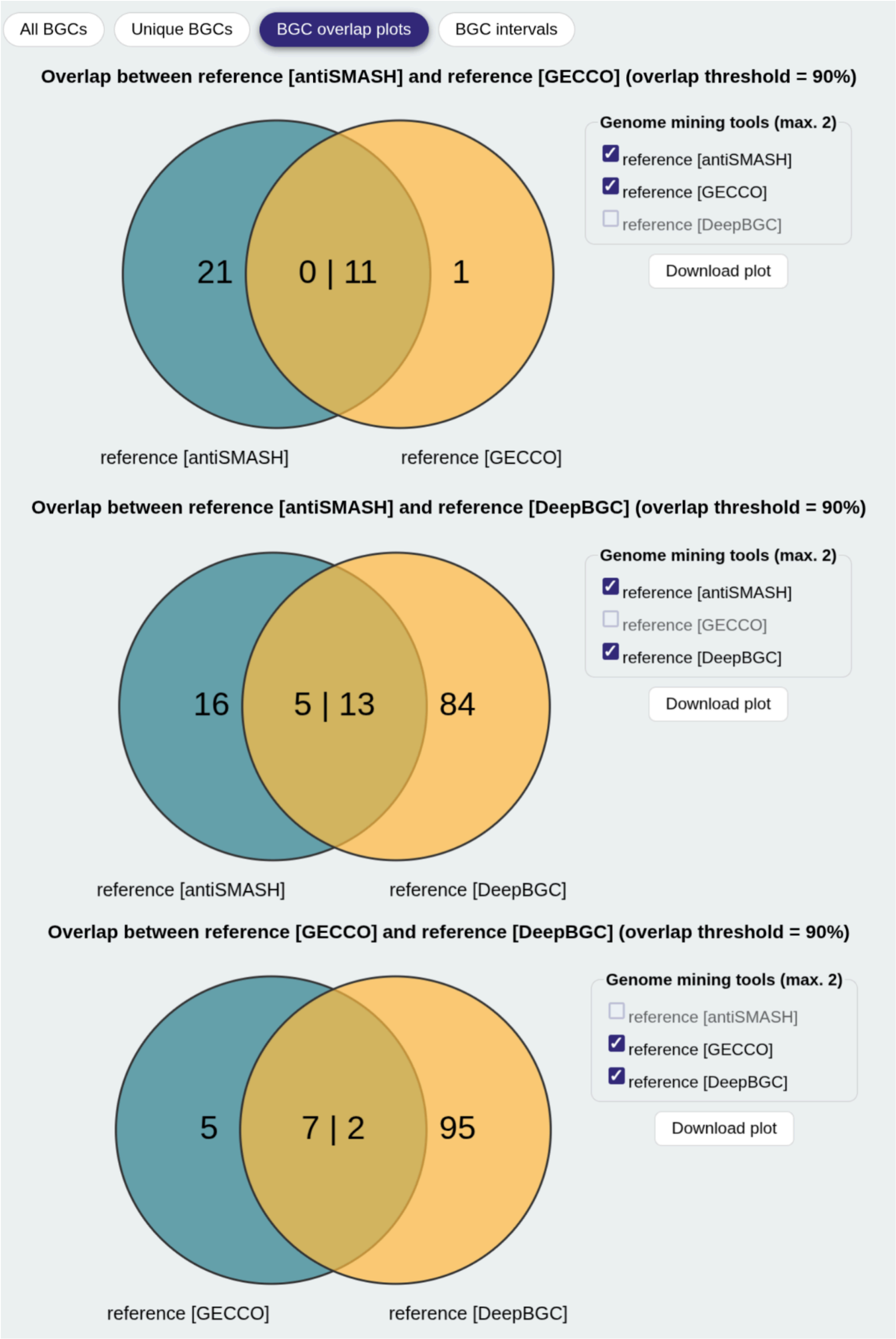


**Supplementary Figure S4.** Examples of BGC overlap plots in a BGC-QUAST compare-tools report. The underlying source data is the same as in Supplementary Figure S3. The three panel switches visualize pairwise overlaps between tool predictions using Venn diagrams, where each diagram shows the number of BGCs unique to each tool and those shared between them. Note that the number of shared BGCs is not necessarily symmetric because the overlap fraction is calculated relative to the length of the BGC being evaluated; therefore, two values are shown in the intersection.

Show 10 rows — Showing 1–10 of 94 Previous 1 / 10 Next

| Sequence ID | Interval start | Interval end | reference<br>antiSMASH | reference<br>GECCO | reference<br>DeepBGC |
|---|---|---|---|---|---|
| Sequence ID | Min | Max | Product | Product | Product |
| 700508.PRJNA177066.CP006936 | 11081 | 107761 | 12076 - 35462 (RiPP);<br>62335 - 107761 (PKS) | 12102 - 30037 (Unknown product);<br>79062 - 93515 (PKS) | 11081 - 104229 (PKS) |
| 700508.PRJNA177066.CP006936 | 145272 | 198061 | 154612 - 198061 (NRPS) | - | 145272 - 163811 (Unknown product);<br>165473 - 166737 (RiPP);<br>174210 - 180402 (NRPS) |
| 700508.PRJNA177066.CP006936 | 204032 | 205078 | - | - | 204032 - 205078 (Unknown product) |
| 700508.PRJNA177066.CP006936 | 218055 | 222004 | - | - | 218055 - 222004 (PKS) |
| 700508.PRJNA177066.CP006936 | 224496 | 224825 | - | - | 224496 - 224825 (Unknown product) |
| 700508.PRJNA177066.CP006936 | 512715 | 518349 | - | - | 512715 - 518349 (Unknown product) |
| 700508.PRJNA177066.CP006936 | 561323 | 584835 | 561323 - 584835 (Terpene) | - | - |
| 700508.PRJNA177066.CP006936 | 592413 | 689442 | 632798 - 683690 (PKS) | 651521 - 675657 (PKS) | 592413 - 671946 (PKS, NRPS);<br>676965 - 689442 (Other) |
| 700508.PRJNA177066.CP006936 | 691021 | 714848 | - | - | 691021 - 714848 (Unknown product) |
| 700508.PRJNA177066.CP006936 | 715756 | 717071 | - | - | 715756 - 717071 (PKS) |

**Supplementary Figure S5.** Example fragment of the interactive BGC intervals table generated in BGC-QUAST compare-tools mode. The underlying source data are the same as in Supplementary Figure S3. BGC predictions from antiSMASH, GECCO, and DeepBGC are grouped into genomic intervals based on coordinate overlap, with coordinates and product types shown for each tool. The table allows filtering by sequence ID, interval coordinates, and product type. Interval grouping is independent of the overlap threshold used to calculate shared and unique BGC metrics in the main compare-tools report.

### Supplementary Note S1. BGC shared/unique classification in the compare-tools mode

In the compare-tools mode, BGC-QUAST determines shared and unique BGCs both pairwise and overall. Internally, each BGC is represented as a 0-based, end-exclusive genomic interval. For an evaluated BGC `bgc_A` predicted by tool `A` and each BGC `bgc_B` predicted by tool `B` on the same sequence record (e.g. contig, scaffold, or chromosome), the overlap fraction is defined as the length of the intersection of their intervals divided by the length of `bgc_A`. Because the overlap fraction is defined relative to the evaluated BGC, the same pair can yield a different overlap value when assessed in the opposite direction. If the overlap fraction for at least one `bgc_B` reaches the `--overlap-fraction` threshold (default 0.9), `bgc_A` is classified as **shared** with respect to tool B; otherwise, it is classified as **unique**.

The core pairwise classification can be summarized in the following pseudocode.

```
function classify_bgc(bgc_A, tool_B_bgcs, threshold=0.9):

    for each bgc_B in tool_B_bgcs on the same sequence record as bgc_A:

        overlap = max(0,
            min(bgc_A.end, bgc_B.end) - max(bgc_A.start, bgc_B.start)
        )

        overlap_fraction = overlap / (bgc_A.end - bgc_A.start)

        if overlap_fraction >= threshold:
            return "shared"

    return "unique"
```

By applying this function to all BGCs predicted by tool `A`, BGC-QUAST determines the number of BGCs from tool `A` that are **shared** with or **unique** with respect to tool `B`. These pairwise counts are used to construct the pairwise Venn diagrams in the compare-tools report. By repeating this comparison against all other tools, BGC-QUAST also determines overall uniqueness for each BGC predicted by tool `A`: a BGC is classified as **unique overall** if it is unique with respect to all other genome mining tools. The resulting counts are reported as the Unique BGCs and Unique rate metrics in the main compare-tools report.

### Supplementary Note S2. Example commands and software versions

#### antiSMASH version 8.0.4

```
antismash <genome.fasta> -c 1 --output-dir <out_dir> --genefinding-tool
prodigal
```

#### GECCO version 0.9.10

```
gecco run --genome <genome.fasta> -o <out_dir>
```

#### DeepBGC version 0.1.31

```
deepbgc pipeline <genome.fasta> -o <out_dir>
```

#### QUAST version 5.3.0

```
quast.py -r reference.fasta Otu207-* -o quast-results
```

#### BGC-QUAST version 1.1.0

```
bgc-quast --mode compare-to-reference -r
genome-mining-results/Otu207/antiSMASH/reference/reference.json
genome-mining-results/Otu207/antiSMASH/Otu207-hybrid/Otu207-hybrid.json
genome-mining-results/Otu207/antiSMASH/Otu207-short/Otu207-short.json -q
genomes/Otu207/quast-results -o bgc-quast-reports/compare-to-reference

bgc-quast --mode compare-tools
genome-mining-results/Otu207/antiSMASH/reference/reference.json
genome-mining-results/Otu207/GECCO/reference/reference.clusters.tsv
genome-mining-results/Otu207/DeepBGC/reference/reference.bgc.tsv -G
genomes/Otu207/reference.fasta -o bgc-quast-reports/compare-tools

cd genome-mining-results/selected-references/antiSMASH/
bgc-quast --mode compare-samples 452863.PRJNA20011/452863.PRJNA20011.json
469371.PRJNA20737/469371.PRJNA20737.json
479435.PRJNA21089/479435.PRJNA21089.json
526225.PRJNA29547/526225.PRJNA29547.json
987045.PRJNA63165/987045.PRJNA63165.json
700508.PRJNA177066/700508.PRJNA177066.json
1246995.PRJNA178199/1246995.PRJNA178199.json
251221.PRJNA9606/251221.PRJNA9606.json
1173027.PRJNA158839/1173027.PRJNA158839.json
290397.PRJNA12634/290397.PRJNA12634.json --names "Actino PRJNA20011","Actino
PRJNA20737","Actino PRJNA21089","Actino PRJNA29547","Actino
PRJNA63165","Actino PRJNA177066","Actino PRJNA178199","Cyano
PRJNA9606","Cyano PRJNA158839","Myxo PRJNA12634" -o
../../../bgc-quast-reports/compare-samples
```
